# ResFinderFG v2.0: a database of antibiotic resistance genes obtained by functional metagenomics

**DOI:** 10.1101/2022.10.19.512667

**Authors:** Rémi Gschwind, Svetlana Ugarcina Perovic, Maja Weiss, Marie Petitjean, Julie Lao, Luis Pedro Coelho, Etienne Ruppé

## Abstract

Metagenomics can be used to monitor the spread of antibiotic resistance genes (ARGs). ARGs found in databases such as ResFinder and CARD primarily originate from culturable and pathogenic bacteria, while ARGs from non-culturable and non-pathogenic bacteria remain understudied. Functional metagenomics is based on phenotypic gene selection and can identify ARGs from non-culturable bacteria with a potentially low identity shared with known ARGs. In 2016, the ResFinderFG v1.0 database was created to collect ARGs from functional metagenomics studies. Here, we present the second version of the database, ResFinderFG v2.0, which is available on the Center of Genomic Epidemiology web server (https://cge.food.dtu.dk/services/ResFinderFG/). It comprises 3,913 ARGs identified by functional metagenomics from 50 carefully curated datasets. We assessed its potential to detect ARGs in comparison to other popular databases in gut, soil and water (marine + freshwater) Global Microbial Gene Catalogues (https://gmgc.embl.de). ResFinderFG v2.0 allowed for the detection of ARGs that were not detected using other databases. These included ARGs conferring resistance to beta-lactams, cycline, phenicol, glycopeptide/cycloserine and trimethoprim/sulfonamide. Thus, ResFinderFG v2.0 can be used to identify ARGs differing from those found in conventional databases and therefore improve the description of resistomes.

**GRAPHICAL ABSTRACT:** 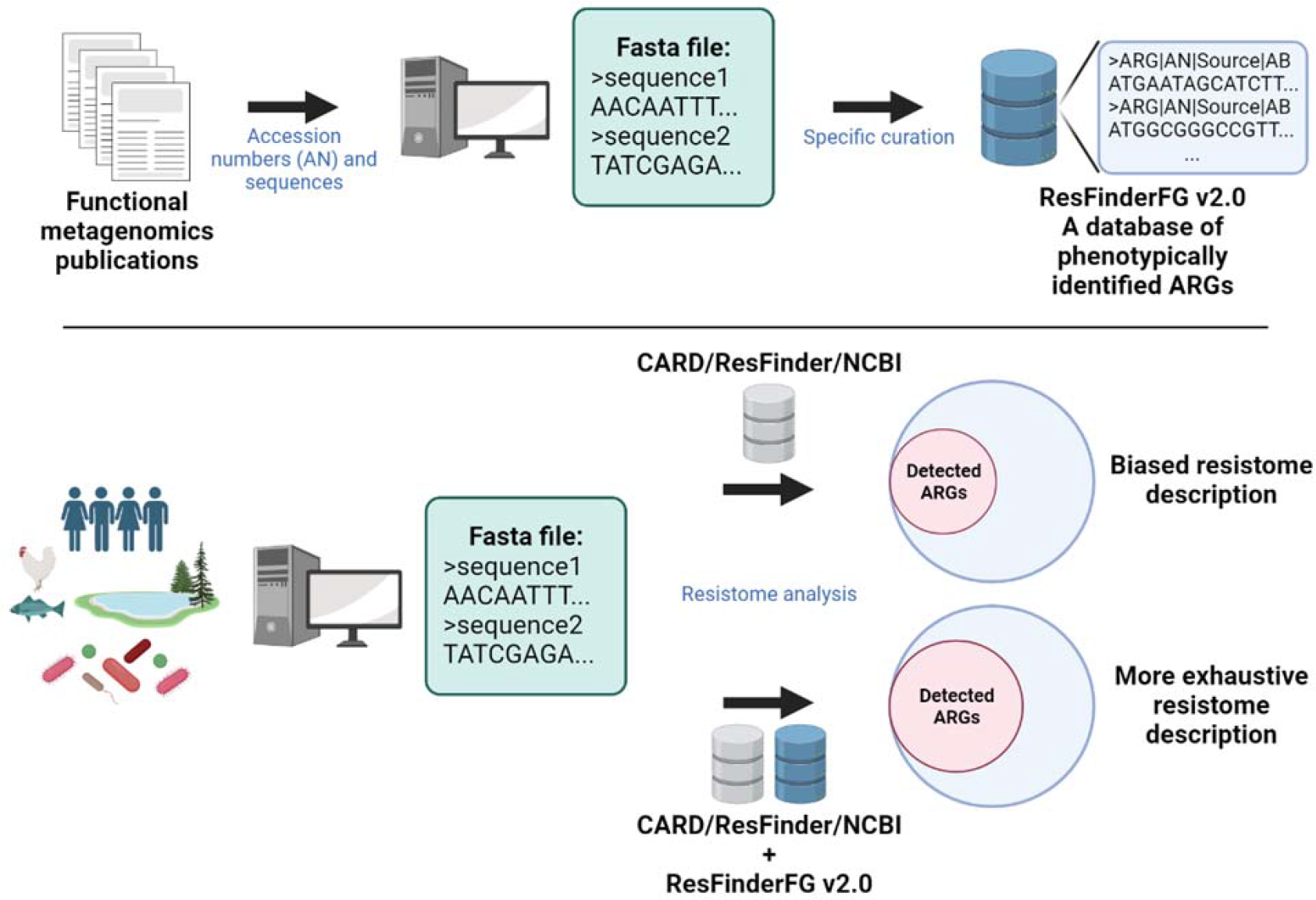

Caption: Additional use of ResFinderFG v2.0 database (composed of antibiotic resistance genes obtained with functional metagenomics) on the Center of Genomic Epidemiology webserver (https://cge.food.dtu.dk/services/ResFinderFG/), allows for more exhaustive resistome descriptions.

## INTRODUCTION

Antimicrobial resistance (AMR) is recognized as a global threat possibly leading to the lack of efficient treatment against deadly infections^1^. From a genetic perspective, AMR is driven by mutational events (e.g. fluoroquinolone resistance is driven by mutations in the topoisomerase-encoding genes) and the expression of antibiotic resistance genes (ARGs). ARGs are widespread in human- and animal-associated microbiomes, and in the environment^2^. Hence, these microbial niches are now considered in a One Health manner^3^. Although not every ARGs represents a direct risk for human health^4^, genes are able to travel from one environment to another by strain dissemination and then to pathogenic bacteria by horizontal gene transfer^5^.

Identifying ARGs and assessing this risk is essential to better understand and putatively find means to prevent their dissemination in pathogenic bacteria. To identify ARGs, culture-based methods, PCR, qPCR^6^, genomic and metagenomic sequencing have been used. Metagenomics makes it possible to sequence all the DNA from a sample. Then with the help of alignment (e.g. BLAST^7^) or Hidden Markov Models (e.g. HMMER^8^) – based tools, ARGs can be identified based on their sequence identity with known ARGs. To this end, several ARG databases^9^ exist such as CARD^10^ or ResFinder^11^. However, their sensitivity and specificity are highly dependent on which database is used and on the parameters used for the search. Moreover, methods using databases of known ARG cannot detect truly novel ARG i.e. not described to date. Pairwise Comparative Modelling (PCM), combining sequence and structure homology to detect ARGs, showed good performances to detect ARGs distantly related to known ARGs in the human gut microbiota, yet some predictions were not all functionally validated^12^.

Although culturable and/or pathogenic bacteria only represent a small fraction of microbial diversity, their genes make up the vast majority of the ARGs present in existing databases. Thus, the description of the resistome can be biased in environments where non-culturable or non-pathogenic bacteria dominate. In order to detect ARGs distantly-related to known ARGs or ARGs from novel ARG families, functional metagenomics has been used^13^. This method is based on phenotypic detection by expressing exogenous DNA in an antibiotic-susceptible host. Metagenomic DNA is sheared and cloned into an expression vector used to transform a host strain susceptible to antibiotics. Transformant strains are then selected with culture media supplemented with antibiotics. If growth is observed, it implies that the DNA cloned into the expression vector is responsible for the resistance phenotype observed. DNA inserts can then be amplified, sequenced and annotated to identify ARGs^14^. Using functional metagenomics, ARGs sharing low amino acid identity to their closest homologue in NCBI^15^, or even not previously classified as ARGs^16^, could be detected in human^16–26^, animal^22,27–33^, wastewater^34–41^ and other environmental samples^5,15,42–64^. Despite being a laborious technique, genes described by functional metagenomics are mainly absent in classical ARG databases. Two databases listing specifically functionally identified ARGs were created: ResFinderFG v1.0^65^ and FARME DB^66^. ResFinderFG v1.0 (https://cge.food.dtu.dk/services/ResFinderFG-1.0/) was based on the data coming from 4 publications, while FARME DB includes data from 30 publications, mainly reporting environmental genes which were not necessarily curated to include only ARG sequences^67^. Here, we report a new version of the ResFinderFG database, ResFinderFG v2.0, providing well-curated data from functional metagenomics publications available until 2021 that include environmental and host-associated samples.

## MATERIAL AND METHODS

### Construction of ResFinderFG v.2.0

To retrieve publications using functional metagenomics for the identification of antibiotic resistance genes, the four publications used to construct ResFinderFG v1.0 were first considered. Then, all the publications which were cited by these four publications and all the publications that cited one of these publications were collected. In addition, publications found with the following terms on PubMed: “functional metagenomics” AND “antibiotic resistance”, were added to this pool. After filtering out all the reviews, publications were screened one by one. First, the use of functional metagenomics to detect ARGs, meaning the expression of exogenous DNA thanks to an expression vector into a host selected on an antibiotic containing medium, was checked. Second, the accessibility to the insert sequences was also checked. Database construction and curation was then performed as follows (Figure 1). The accession numbers describing insert DNA sequences functionally selected using antibiotics were included and DNA sequences were retrieved using Batch Entrez. CD-HIT^68^ was used to remove redundant DNA sequences and annotation of the remaining was done using PROKKA v.1.14^69^. To specifically select insert DNA sequences with ARG annotations, a representative pool of ARG annotations was obtained by applying the PROKKA annotation process to the ResFinder v4.0 database. Resulting annotations were used as a reference to specifically select insert DNA sequences containing an ARG annotation. Metadata (sample origin and antibiotic used for selection) associated with each insert were collected. Then, several curation steps were added. First, a curation based on the consistency between the ARG and the antibiotic family used for selection (e.g. beta-lactamase-encoding gene selected with a beta-lactam antibiotic). Second, we apply a filter on the size in amino acids of the predicted protein: 260 for beta-lactamase, 378 for tetracycline efflux genes, 641 for tetracycline resistance ribosomal protection genes, 178 for chloramphenicol acetyltransferase, 247 for methyltransferase genes and 158 aa for dihydrofolate reductase genes). Finally, insert sequences containing more than one ARG annotation (consistent with the antibiotic used for selection) were discarded as we could not know which one was responsible for the observed phenotype. The database also includes metadata retrieved in the GenBank metadata retrieval process and ARO annotation for each gene for comparability with other databases using ARO ontologies^10^. Besides, we provide in the supplementary material the access numbers of inserts (n=5’256) containing a single ORF among which potential ARG from new families can be characterized (Supplementary Table 1).

**Figure 1:**
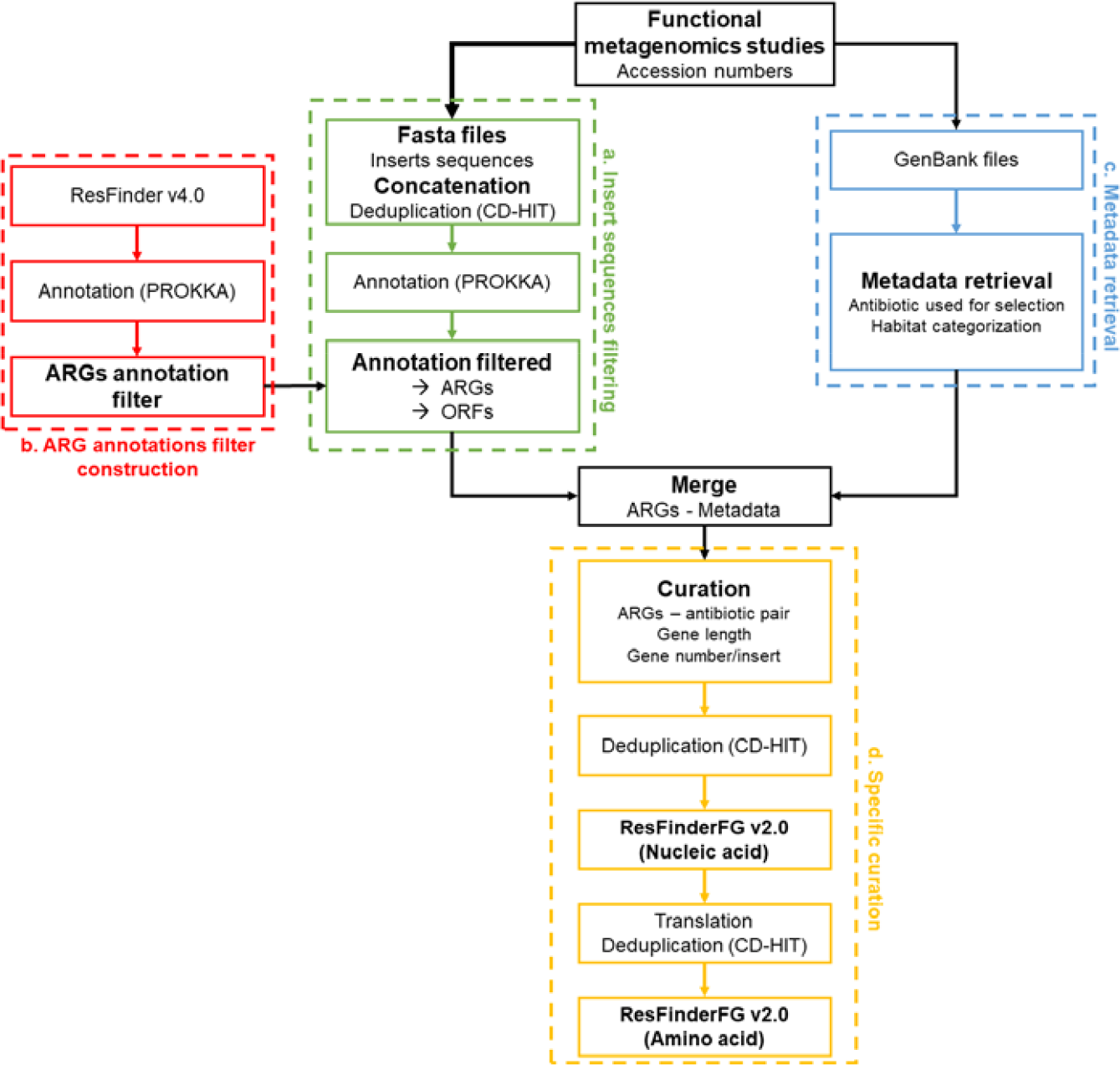
ResFinderFG v.2.0 construction workflow. a. The insert sequences filtering step consists of concatenating all the insert sequences and selecting only those containing an ARG annotation. b. ARG annotations filter construction was done by annotating the ResFinder v4.0 database using PROKKA to have a pool of ARG annotations. c. Metadata was retrieved using GenBank files to get the origin of the samples and the antibiotic used for selection of each insert sequence. d. Specific curation was done to discard false positives and include only ARGs.

### Comparison between ResFinderFG v1.0 and ResFinderFG v2.0

To assess the update of the ResFinderFG database and compare v2.0 to v1.0, the number of ARGs in each ARG family and regarding the sample sources was evaluated in both versions. ARG families were categorized according to the antibiotic families they conferred resistance to: glycopeptides/cycloserine, sulfonamides/trimethoprim, beta-lactams, aminoglycosides, macrolides/lincosamides/streptogramins, tetracyclines, phenicols and quinolones. Sample sources were categorized as follows: aquatic, animal-associated, human-associated, plant-associated, polluted environment and soil. Then, to detect the presence of ARGs in several gene subcatalogs (human gut, soil and marine+freshwater) coming from the Global Microbial Gene Catalog (GMGC, https://gmgc.embl.de/download.cgi^2^), ABRicate^70^ was run using default parameters with different databases (ResFinderFG v2.0, ResFinder v4.0, CARD v3.0.8, ARG-ANNOT v5, NCBI v3.6).

## RESULTS

### Construction of ResFinderFG v2.0

A total of 50 publications using functional metagenomics to analyse ARG content were selected, resulting in 23,776 accession numbers (Supplementary Table 2). CD-HIT identified 2,629 perfectly redundant insert sequences (100% sequence identity). PROKKA identified 41,977 open reading frames (ORFs). Among them, 7,787 ORFs had an annotation matching with an ARG annotation of ResFinder v4.0 (228 unique ARG annotations). Then, we did not consider the following ARGs: (i) inconsistency between the predicted ARG and the antibiotic used for selection (n=1,165), (ii) ARGs with excessively small size with respect to the ARG family (n=1,064) and ARGs colocating on an insert (n=398). A second round of CD-HIT was used to remove redundancy (100% nucleic acid sequence identity), which resulted in 3,913 unique ARGs which were finally included in the database which can be used on the Center of Genomic Epidemiology website (https://cge.food.dtu.dk/services/ResFinderFG/; supplementary figure 1).

### Comparison with ResFinderFG v1.0

First, the ARGs present in ResFinderFG v.2.0 were compared to the ones present in ResFinderFG v1.0 (Figure 2). A total of 1,631 new ARGs were present in ResFinderFG v.2.0, mainly due to new glycopeptide/cycloserine (+906 genes) and beta-lactam (+333 genes) resistance genes. The glycopeptide/cycloserine resistance genes were mostly annotated as homologues of D-Ala-D-X ligase. New beta-lactams antibiotics used for functional selection compared to v1.0 were cefepime, meropenem and tazobactam. Regarding the sources of ARGs, new ARGs mostly originated from human-associated samples (+1,333 genes).

**Figure 2:**
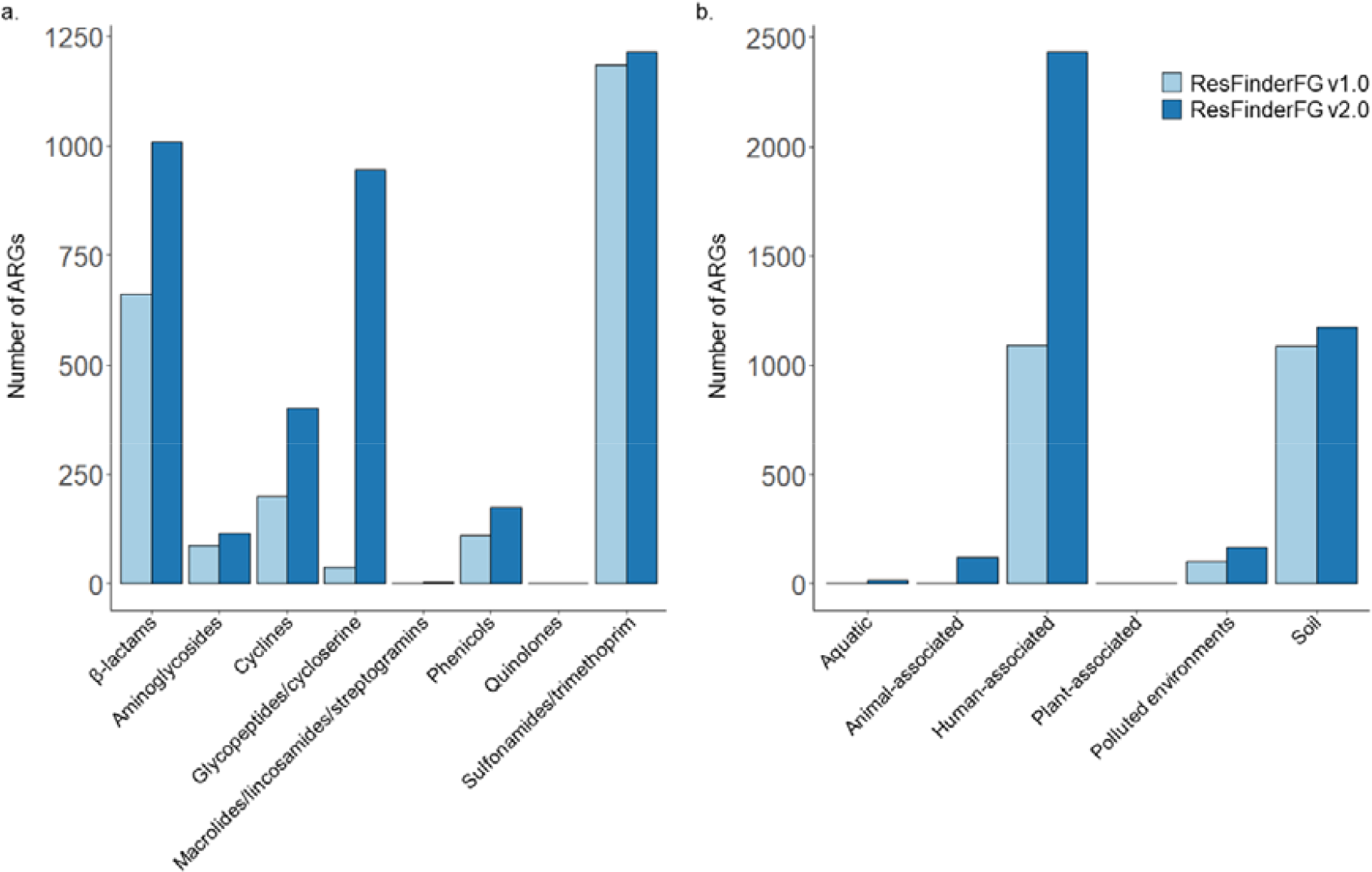
a. Number of ARGs in the ResFinderFG v.1.0 and v.2.0 databases depending on a. the antibiotic families involved; b. the sample sources.

### ARG detection in several GMGC gene subcatalogs using ResFinderFG v.2.0 and other databases

ABRicate (default parameters) was used to detect ARGs in GMGC human gut (Figure 3), soil (Supplementary Figure 2a.) and aquatic (marine and freshwater) subcatalogs (Supplementary Figure 2b.). Using ResFinderFG v2.0, 3,025, 211 and 129 unigene hits were obtained analyzing human gut, soil and aquatic subcatalogs respectively. The three most frequently detected ARG families in all gene catalogs were glycopeptide/cycloserine resistance genes (20.9 to 39.7% of detected ARGs), sulfonamides/trimethoprim resistance genes (21.8 to 58.1% of detected ARGs) and beta-lactamase encoding genes (7.9 to 25.6% of detected ARGs). Phenicols (up to 6.0% of detected ARGs), aminoglycosides (up to 5.3%), cyclines (up to 6.2%) and macrolide/ lincosamide/streptogramin resistance genes (up to 0.03%) were also detected. Also, ResFinderFG v2.0 provides habitat information on where a given ARG was first identified by functional metagenomics. A majority of ARGs identified in the gut subcatalog (90.2%) were indeed initially identified in the human gut by functional metagenomics (supplementary table 3). In the soil gene subcatalog, 62.6% of ARGs detected were also genes identified initially in soil with functional metagenomics. However, ARGs detected in the aquatic gene subcatalog were primarily first identified by functional metagenomics in soil.

**Figure 3.**
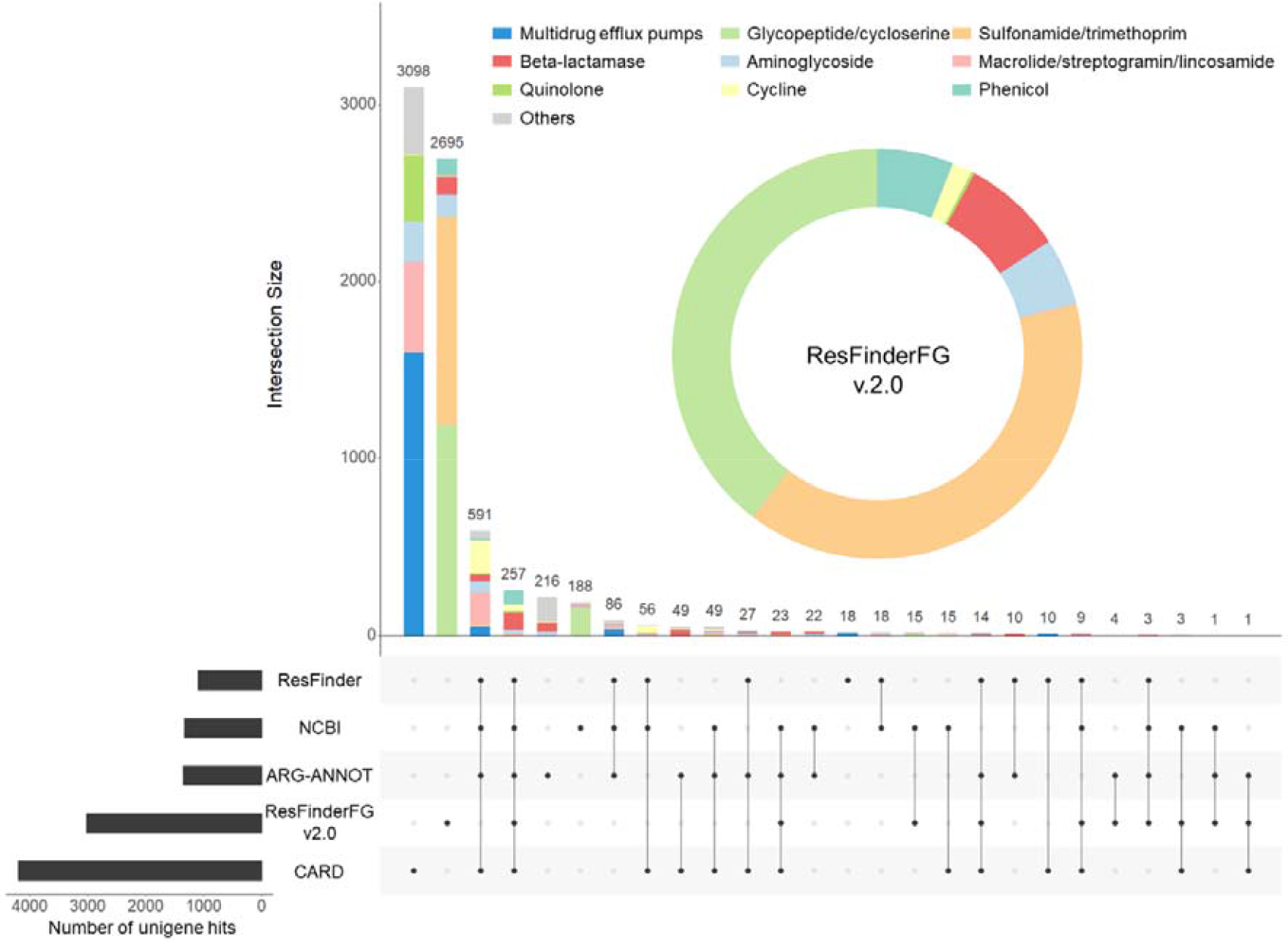
Proportion of antibiotic family unigene hits obtained analyzing GMGC human gut subcatalog using ResFinderFG v2.0 and number of unigene hits obtained analyzing GMGC human gut subcatalog using several databases (ResFinder v4.0, NCBI v3.6, ARG-ANNOT v5, ResFinderFG v2.0 and CARD v3.0.8) annotated by their antibiotic family. Others: bicyclomycin, beta-lactams, bleomycin, disinfectant and antiseptic agents, fosfomycin, fusidic acid, multidrug, mupirocin, nitroimidazole, nucleoside, peptide, rifampicin, streptothricin.

To compare ResFinderFG v2.0 to other databases, we ran the same ABRicate analysis of GMGC gene subcatalogs using ResFinder v.4.0, CARD v3.0.8, ARG-ANNOT v5 and NCBI v3.6. ResFinderFG v2.0 identified a comparable or even greater number of ARGs compared to other databases. We observed that the most frequently observed ARG family depended on the database used. In the human gut gene subcatalog, glycopeptide/cycloserine resistance gene was the most frequent ARG family found by ResFinderFG v2.0 (39.7% of all unigene hits obtained with ResFinderFG v2.0). In contrast, the beta-lactamase family was the top ARG family with ARG-ANNOT (21.2%). NCBI and ResFinder detected mostly tetracycline resistance genes (20.4 and 23.8% respectively). Finally, multidrug efflux pump unigene hits were the most frequent using CARD (39.4%).

ResFinderFG v2.0 was the database with the highest fraction of database-specific hits, with 89.1% of specific unigene hits composed mainly by glycopeptide/cycloserine resistance genes (D-alanine-D-alanine ligase; supplementary table 4) and sulfonamides/trimethoprim resistance genes (dihydrofolate reductase). By comparison, CARD had 73.7% of specific unigene hits, mostly composed by gene encoding multidrug efflux pumps. Of note, 16.2% of unique CARD specific multidrug efflux pump unigene hits found in the human gut were regulatory genes (Supplementary Table 4).

Between 2.6 and 4.2% of all unigene hits, depending on the gene subcatalog analyzed, were shared by all the databases used. Beta-lactamase encoding genes were the most prevalent among them (ranging from 38.1 to 51.3% of the shared unigene hits), followed by, phenicols, aminoglycosides and tetracyclines resistance genes. However, 25.1, 23.2 and 46.3% of beta-lactamases, aminoglycosides and phenicols resistance genes respectively, were only detected using ResFinderFG v2.0 (Figure 3; supplementary Figure 2).

## DISCUSSION

ResFinderFG v2.0 contains 3,913 ARGs which were described with functional metagenomics in 50 publications. Here, we showed that using ResFinderFG v2.0 enabled us to describe the resistome with ARGs that were not detected by other databases. Notably, ResFinderFG v2.0 permitted a better description of sulfonamide/trimethoprim, glycopeptide/cycloserine resistant genes and beta-lactamase encoding genes.

ResFinderFG v2.0 includes more ARGs coming from human-associated samples^16–26^. For example, characterization of the gut resistome with functional metagenomics showed that its ARGs were not well described in ARG databases^18^. Inclusion of these ARGs is therefore important for future metagenomic characterization of resistomes. Regarding the ARG family concerned, most of the new ARGs included compared to ResFinderFG v1.0 are glycopeptide/cycloserine or beta-lactam resistance genes. Glycopeptide/cycloserine resistance genes were selected using cycloserine, an antibiotic used in the therapy of tuberculosis caused by multi resistant mycobacteria^71^. Beta-lactam resistant genes are of high concern because beta-lactam antibiotics are widely used against priority pathogens^72^.

Using ResFinderFG v2.0, sulfonamide/trimethoprim, glycopeptide/cycloserine, beta-lactam, phenicol, cycline, quinolone, macrolide/lincosamide/streptogramin and aminoglycoside resistance genes were evidenced studying three GMGC gene subcatalogs (human gut, soil and aquatic). As expected, regarding their representation in the database, the most frequent unigene hits were glycopeptide/cycloserine and sulfonamide/trimethoprim resistance genes. Analogous analyses performed with other databases showed that ResFinderFG v2.0 detected a comparable or higher number of ARGs depending on the other database used. Beta-lactamase encoding genes were the most represented ARGs in unigene hits shared by all databases. Yet, ResFinderFG v2.0 allowed the detection of beta-lactamase encoding genes which were not detected with other databases. It was expected since many publications using functional metagenomics reported beta-lactamase encoding genes distant from the ones described in ARG databases^15,18,35,38,50,58,63^ and a distant one has been evidenced recently from soil samples^43^. Other antibiotic families were even more specifically associated with ResFinderFG v2.0, such as sulfonamide/trimethoprim, phenicol and glycopeptide/cycloserine resistance genes.

Our study has limitations, though. The strength of functional metagenomics is its ability to identify ARGs without relying on their DNA sequence and to provide phenotypical data associated with ARGs. This makes it possible to identify new resistance mechanisms from new families of ARGs. However, to properly identify the latter and avoid false positives, substantial laboratory work is required and is not done for most inserts in functional metagenomics experiments. In order to characterise an ARG, further cloning is required to study which part of the insert is important for the observation of the phenotype and also to what extent this may modify the associated MICs. Consequently, ResFinderFG v2.0 includes ARGs belonging to known families as opposed to truly novel ARGs from new families which would require further experiments to be identified. We are also aware that some genes included in ResFinderFG v2.0 require overexpression to confer resistance to an antibiotic. This is the case for example for D-alanine-D-alanine ligase or dihydrofolate reductase encoding genes which confer resistance to cycloserine or trimethoprim respectively when they are overexpressed. Yet, these genes are also described in other ARG databases and they fit the operational definition of an ARG^4^. Finally, since the original accession numbers are available in each ARG sequence header, researchers can easily obtain the complete insert DNA sequence to investigate.

## DATA AVAILABILITY

All the computational steps and data used in the construction of the ResFinderFG v2.0 database and the database itself are available on the following public repository: https://zenodo.org/badge/latestdoi/470536952. The database was also deposited on the Center of Genomic Epidemiology (CGE) web server, where it can be used online https://cge.food.dtu.dk/services/ResFinderFG/. Supplementary figure 1 describes how to use the web server. Analysis processes for the description of ResFinderFG v2.0 are accessible on the following public repository: https://zenodo.org/badge/latestdoi/507027650.

## SUPPLEMENTARY DATA

Supplementary Data are available at NAR online.

## AUTHOR CONTRIBUTIONS

RG made the database update, the comparison analysis of ABRicate results with other databases and wrote the manuscript. SUP made the initial ABRicate analysis on GMGC subcatalogs and reviewed the manuscript. JL and MP helped building the figures and the scripts for database construction and analysis. MW made the update available on the CGE webserver. LPC reviewed the manuscript. ER designed the study and reviewed the manuscript.

## ACKNOWLEDGEMENTS

The authors are grateful to Frank Møller Aarestrup for hosting ResFinderFG v2.0 on the Center for Genomic Epidemiology website, and to Andrew Bielski for English editing. Graphical abstract was created using BioRender.

## FUNDING

This work was funded by the Joint Program Initiative for Antimicrobial Resistance (JPIAMR) EMBARK (Establishing a Monitoring Baseline for Antimicrobial Resistance in Key environments) project (International Development Research Centre, IDRC, grant 109304-001 to LPC, Agence Nationale de la Recherche, ANR, grant ANR-19-JAMR-0004 to ER).

## CONFLICT OF INTEREST

All authors: none

## Notes

### Competing Interest Statement

The authors have declared no competing interest.

### Summary of Updates

We clarified the purpose of our manuscript. We added a new table with accession numbers of insert sequences containing one unique ORF and which could be potential ARGs to be functionally confirmed. We added a new supplementary figure to describe how to use the ResFinderFG v2.0 database on the CGE website.

