## Supplementary materials for "ResFinderFG v2.0: a database of antibiotic resistance genes obtained by functional metagenomics"

**SUPPLEMENTAL DATA**

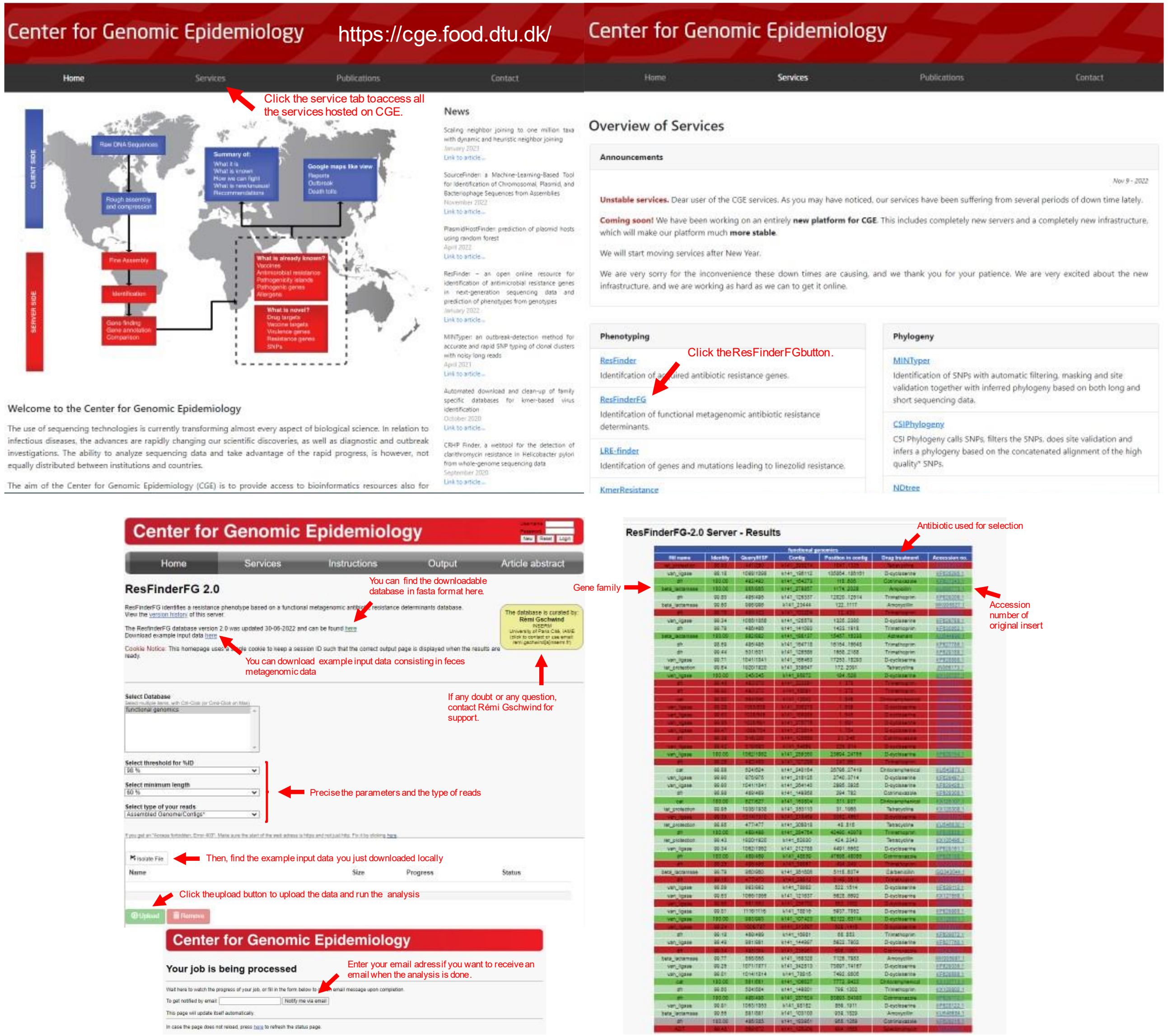


**Supp figure 1:** Instruction for use ResFinderFG v2.0 database on the center of genomic epidemiology website.

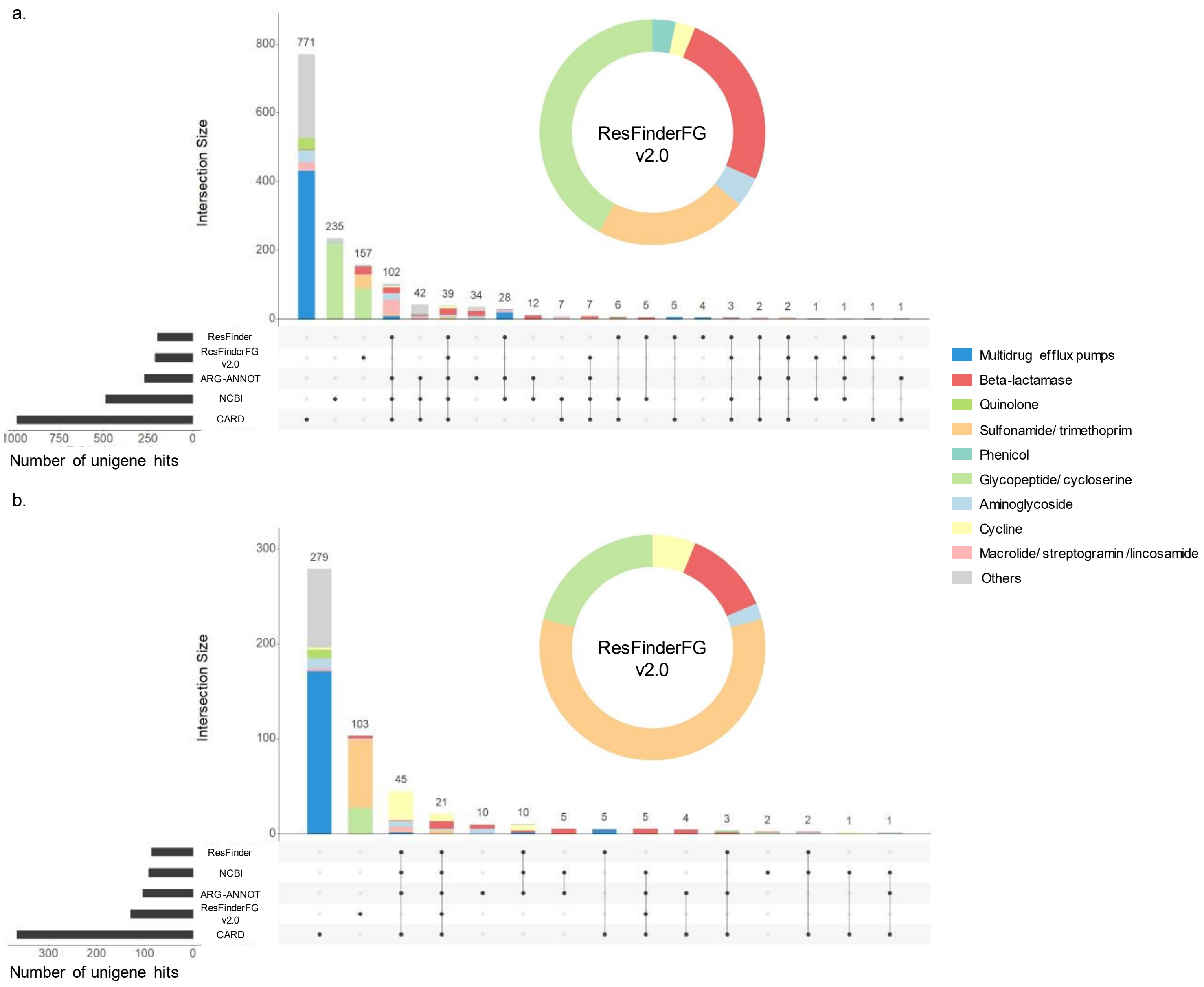
a.
Intersection Size
800
600
400
200
0
771
235
157
102
42
39
34
28
12
7
7
6
5
5
4
3
2
2
1
1
1
1
ResFinderFG
v2.0
ResFinder
ResFinderFG
v2.0
ARG-ANNOT
NCBI
CARD
1000
750
500
250
0
Number of unigene hits
b.
Intersection Size
300
200
100
0
279
103
45
21
10
10
5
5
5
4
3
2
2
1
1
ResFinderFG
v2.0
ResFinder
NCBI
ARG-ANNOT
ResFinderFG
v2.0
CARD
300
200
100
0
Number of unigene hits
Multidrug efflux pumps
Beta-lactamase
Quinolone
Sulfonamide/ trimethoprim
Phenicol
Glycopeptide/ cycloserine
Aminoglycoside
Cycline
Macrolide/ streptogramin /lincosamide
Others

**Supp Figure 2:** Proportion of antibiotic family unigene hits obtained analyzing GMGC a. soil and b. aquatic subcatalog (freshwater and marine) and number of ARGs detected using several databases (ResFinder v4.0, NCBI v3.6, ARG-ANNOT v5, ResFinderFG v2.0 and CARD v3.0.8) regarding the antibiotic family concerned. Others: bicyclomycin, beta-lactams, bleomycin, disinfectant and antiseptic agents, fosfomycin, fusidic acid, multidrug, mupirocin, nitroimidazole, nucleoside, peptide, rifampicin, streptothricin.

**Supp Table 1:** Inserts including 1 ORF but which were excluded from ResFinderFG v2.0 for: (i) annotation not corresponding to an ARG annotation; (ii) unconsistency between antibiotic used for selection and ARG family; (iii) irrelevant gene size.

**Supp Table 2:** Publications included in ResFinderFG v2.0 construction.

| Year | Author | Title | Accession numbers | Insert sequences | Deduplicated insert sequences | ORFs | doi |
|---|---|---|---|---|---|---|---|
| 2021 | McGivern | Novel class 1 integron harboring antibiotic resistance genes in wastewater-derived bacteria as revealed by functional metagenomics | MN340011-MN340023 | 13 | 13 | 503 | 10.1016/j.plasmid.2021.102563 |
| 2021 | Willms | Novel soil-derived beta-lactam, chloramphenicol, fosfomycin and trimethoprim resistance genes revealed by functional metagenomics | MW601939-MW601946 | 8 | 8 | 39 | 10.3390/antibiotics10040378 |
| 2021 | Obermeïer | Plant resistome profiling in evolutionary old bog vegetation provides new clues to understand emergence of multi-resistance | MK831000 | 1 | 1 | 1 | 10.1038/s41396-020-00822-9 |
| 2020 | Reynolds | Detection of a Novel, and Likely Ancestral, Tn *916*-Like Element from a Human Saliva Metagenomic Library | MF344584 | 1 | 1 | 12 | 10.3390/genes11050548 |
| 2020 | Rahman | Integron gene cassettes harboring novel variants of D-alanine-D-alanine ligase confer high-level resistance to D-cycloserine | KU886208-KU886209 | 2 | 2 | 4 | 10.1038/s41598-020-77377-4 |
| 2020 | Wang | Duck wastes as a potential reservoir of novel antibiotic resistance genes | MW234453-MW234459 | 2 | 2 | 2 | 10.1016/j.scitotenv.2020.144828 |
| 2020 | Böhm | Discovery of a novel integron borne aminoglycoside resistance gene present in clinical pathogens by screening enviornmental bacterial communities | MN215968.1 | 1 | 1 | 1 | 10.1186/s40168-020-00814-z |
| 2020 | Campbell | The microbiome and resistome of chimpanzees, gorillas, and humans across host lifestyle and geography | MK935708-MK936039 | 332 | 218 | 207 | 10.1038/s41396-020-0634-2 |
| 2019 | Marathe | Sewage effluent from an Indian hospital harbors novel carbapenemases and integron-borne antibiotic resistance genes | MN017279-MN017299 | 21 | 21 | 21 | 10.1186/s40168-019-0710-x |
| 2019 | Zhang | Novel clinically relevant antibiotic resistance genes associated with sewage sludge and industrial waste streams revealed by functional metagenomic screening | KF485391-KF485396 | 6 | 6 | 39 | 10.1016/j.envint.2019.105120 |
| 2019 | Willms | Discovery of Novel Antibiotic Resistance Determinants in Forest and Grassland Soil Metagenomes | MK159018-MK159025 | 8 | 8 | 43 | 10.3389/fmicb.2019.00460 |
| 2018 | Park | PNGM-1, a novel subclass B3 metallo-β-lactamase from a deep-sea sediment metagenome | MF445022 | 1 | 1 | 1 | 10.1016/j.jgar.2018.05.021 |
| 2018 | Marathe | Functional metagenomics reveals a novel carbapenem-hydrolyzing mobile beta-lactamase from Indian river sediments contaminated with antibiotic production waste | MG739504-MG739510 | 7 | 7 | 7 | 10.1016/j.envint.2017.12.036 |
| 2018 | González-Plaza | Functional Repertoire of Antibiotic Resistance Genes in Antibiotic Manufacturing Effluents and Receiving Freshwater Sediments. | MG585943-MG586044 | 102 | 63 | 63 | 10.3389/fmicb.2017.02675 |
| 2017 | Wang | Tetracycline Resistance Genes Identified from Distinct Soil Environments in China by Functional Metagenomics. | KX161706-KX161713<br>KY697282-KY853666 | 24 | 22 | 23 | 10.3389/fmicb.2017.01406 |
| 2017 | Ho-Fung Lau | Novel Antibiotic Resistance Determinants from Agricultural Soil Exposed to Antibiotics Widely Used in Human Medicine and Animal Farming | KY705323-KY705356 | 34 | 34 | 1108 | 10.1128/AEM.00989-17 |
| 2017 | Florez | A Functional Metagenomic Analysis of Tetracycline Resistance in Cheese Bacteria | KY686299-KY686304 | 6 | 6 | 212 | 10.3389/fmicb.2017.00907 |
| 2016 | Pawlowski | A diverse intrinsic antibiotic resistome from a cave bacterium | KX531043-KX531056 | 14 | 14 | 14 | 10.1038/ncomms13803 |
| 2016 | Versluis | Sponge Microbiota Are a Reservoir of Functional Antibiotic Resistance Genes | KU577908-KU577944 | 37 | 37 | 149 | 10.3389/fmicb.2016.01848 |
| 2016 | Gibson | Developmental dynamics of the preterm infant gut microbiota and antibiotic resistome | KU605810-KU608292 | 2483 | 2000 | 4016 | 10.1038/nmicrobiol.2016.24 |
| 2016 | Gudeta | Expanding the Repertoire of Carbapenem-Hydrolyzing Metallo-ß-Lactamases by Functional Metagenomic Analysis of Soil Microbiota | KU167035-KU167043 | 9 | 9 | 9 | 10.3389/fmicb.2016.01985 |
| 2016 | Pehrsson | Interconnected microbiomes and resistomes in low-income human habitats | KU543693-KU549046 | 5354 | 4430 | 7290 | 10.1038/nature17672 |
| 2015 | Clemente | The microbiome of uncontacted Amerindians | KJ910906-KJ911011 | 106 | 96 | 325 | 10.1126/sciadv.1500183 |
| 2015 | Munck | Limited dissemination of the wastewater treatment plant core resistome | KT387137-KT387215 | 79 | 78 | 78 | 10.1038/ncomms9452 |
| 2015 | Hatosy | The ocean as a global reservoir of antibiotic resistance genes | KS307228-KS308058 | 831 | 796 | 699 | 10.1128/AEM.00736-15 |
| 2015 | Perron | Functional Characterization of Bacteria Isolated from Ancient Arctic Soil Exposes Diverse Resistance Mechanisms to Modern Antibiotics | KC520483-KC520492 | 10 | 10 | 10 | 10.1371/journal.pone.0069533 |
| 2015 | Moore | Gut resistome development in healthy twin pairs in the first year of life | KX125104-KX128902 | 3799 | 3473 | 6729 | 10.1186/s40168-015-0090-9 |
| 2014 | Udikovic Kolic | Bloom of resident antibiotic-resistant bacteria in soil following manure fertilization | KM113767-KM113773 | 7 | 7 | 13 | 10.1073/pnas.1409836111 |
| 2014 | Su | Functional metagenomic characterization of antibiotic resistance genes in agricultural soils from China. | JX875536-JX875591 | 56 | 55 | 103 | 10.1016/j.envint.2013.12.010 |
| 2014 | Wichmann | Diverse Antibiotic Resistance Genes in Dairy Cow Manure | KJ512900-KJ512991 | 92 | 80 | 302 | 10.1128/mBio.01017-13 |
| 2014 | Forsberg | Bacterial phylogeny structures soil resistomes across habitats. | KJ691878-KJ696532 | 4655 | 4552 | 10028 | 10.1038/nature13377 |
| 2013 | Gong Cheng | Identification of a novel fosfomycin-resistant UDP-N-acetylglucosamine enolpyruvyl transferase (MurA) from a soil metagenome | JN629036 | 1 | 1 | 1 | 10.1007/s10529-012-1074-5 |
| 2013 | López-Pérez | Identification and modeling of a novel chloramphenicol resistance protein detected by functional metagenomics in a wetland of Lerma, Mexico. | KF169941 | 1 | 1 | 1 | 10.2436/20.1501.01.185 |
| 2013 | Moore | Pediatric Fecal Microbiota Harbor Diverse and Novel Antibiotic Resistance Genes | KF626669-KF630360 | 3692 | 3457 | 7201 | 10.1371/journal.pone.0078822 |
| 2013 | Vercammen | Identification of a metagenomic gene cluster containing a new class A beta-lactamase and toxin-antitoxin systems | KF033132 | 1 | 1 | 38 | 10.1002/mbo3.104 |
| 2013 | Berman | Identification of novel antimicrobial resistance genes from microbiota on retail spinach | KF791056-KF791060 | 5 | 5 | 13 | 10.1186/1471-2180-13-272 |
| 2012 | Zhou | Functional Cloning and Characterization of Antibiotic Resistance Genes from the Chicken Gut Microbiome | JN625754-JN625767 | 14 | 13 | 12 | 10.1128/AEM.06920-11 |
| 2012 | Forsberg | The shared antibiotic resistome of soil bacteria and human pathogens. | JX009202-JX009380 | 179 | 141 | 161 | 10.1126/science.1220761 |
| 2012 | Cheng | Functional screening of antibiotic resistance genes from human gut microbiota reveals a novel gene fusion | JN086157-JN086173 | 17 | 17 | 49 | 10.1111/j.1574-6968.2012.02647.x |
| 2012 | Tian | Long-term exposure to antibiotics has caused accumulation of resistance determinants in the gut microbiota of honeybees | JQ966977-JQ966984<br>JS807327-JS807645 | 327 | 307 | 491 | 10.1128/mBio.00377-12 |
| 2011 | Uyaguari | Characterization and quantitation of a novel β-lactamase gene found in a wastewater treatment facility and the surrounding coastal ecosystem | HQ605913 | 1 | 1 | 1 | 10.1128/AEM.02732-10 |
| 2011 | McGarvey | Wide Variation in Antibiotic Resistance Proteins Identified by Functional Metagenomic Screening of a Soil DNA Library | JF924866-JF924904 | 39 | 39 | 115 | 10.1128/AEM.06759-11 |
| 2011 | Martiny | Functional metagenomics reveals previously unrecognized diversity of antibiotic resistance genes in gulls | JM426721-JM426844 | 1124 | 1102 | 1127 | 10.3389/fmicb.2011.00238 |
| 2011 | Torres-Cortès | Characterization of novel antibiotic resistance genes identified by functional metagenomics on soil samples | FN640464-FN640474 | 11 | 11 | 45 | 10.1111/j.1462-2920.2010.02422.x |
| 2010 | Parsley | Identification of Diverse Antimicrobial Resistance Determinants Carried on Bacterial, Plasmid, or Viral Metagenomes from an activated Sludge Microbial Assemblage | GU720994-GU721005 | 12 | 12 | 12 | 10.1128/AEM.03080-09 |
| 2009 | Allen | Resident Microbiota of the Gypsy Moth Midgut Harbors Antibiotic Resistance Determinants | EU408346-EU408359 | 3 | 3 | 13 | 10.1089/dna.2008.0812 |
| 2009 | Donato | Metagenomic Analysis of Apple Orchard Soil Reveals Antibiotic Resistance Genes Encoding Predicted Bifunctional Proteins | GQ244488-GQ244501 | 14 | 14 | 61 | 10.1128/AEM.01763-09 |
| 2009 | Allen | Functional metagenomics reveals diverse b-lactamases in a remote Alaskan soil | EU885953-EU885955 | 14 | 14 | 314 | 10.1038/ismej.2008.86 |
| 2009 | Sommer | Functional Characterization of the Antibiotic Resistance Reservoir in the Human Microflora | GQ342978-GQ343187 | 210 | 182 | 319 | 10.1126/science.1176950 |
| 2004 | Riesenfeld | Uncultured soil bacteria are a reservoir of new antibiotic resistance genes | AY566820-AY566829 | 10 | 10 | 21 | 10.1111/j.1462-2920.2004.00664.x |

**Supp Table 3:** Unigene hits obtained using ResFinderFG v2.0 in GMGC subcatalogs and the initial habitat associated found in ResFinderFG v2.0 metadata.

| Subcatalog | | total | human gut | soil | other |
|---|---|---|---|---|---|
| Gut | n gene | 3025 | 2729 | 61 | 234 |
| | % | | 90,2 | 2,0 | 7,7 |
| Soil | n gene | 211 | 42 | 132 | 37 |
| | % | | 19,9 | 62,6 | 17,5 |
| water | n gene | 129 | 43 | 82 | 4 |
| | % | | 33,3 | 63,6 | 3,1 |

**Supp Table 4:** Specific genes found in each catalog analysed regarding the database used.

| Database | Antibiotic family | Gut | Soil | Water |
|---|---|---|---|---|
| ARG-ANNOT | Aminoglycoside | (AGly)aadA6; (AGly)aadA4; (AGly)aadC; (AGly)apH-Stph; (AGly)aph7 | (AGly)aadC; (AGly)apH-Stph | (AGly)aadC |
| | non-enzymatic β-lactam resistance | (Bla)Penicillin_Binding_Protein_Ecoli; (Bla)PBP1a; (Bla)PBP1b | (Bla)Penicillin_Binding_Protein_Ecoli | (Bla)Penicillin_Binding_Protein_Ecoli |
| | β-lactamase | (Bla)BlaA1; (Bla)ampH; (Bla)BlaA2; (Bla)ampS; (Bla)Mbl; (Bla)Zn-dependent_hydrolase | (Bla)ampH; (Bla)BlaA1; (Bla)BlaA2; (Bla)blaLMB-1; (Bla)Mbl; (Bla)Zn-dependent_hydrolase | (Bla)Mbl; (Bla)Zn-dependent_hydrolase |
| | cycline | (Tet)tetR(G) | (Tet)tetR(G) | - |
| | MLS | (MLS)mph(D) | - | - |
| | phenicol | (Phe)catB4; (Phe)dha1 | - | - |
| CARD | aminocoumarin | - | point mutation on parY; A type III ABC transporter | - |
| | aminoglycoside | AcrD; BaeR; BaeS; CpxA; EmrE; kdpE | AcrD; BaeR; CpxA | BaeS; AcrD; kdpE; BaeR; CpxA |
| | non-enzymatic β-lactam resistance | OmpK | - | - |
| | β-lactamase | - | NmcR | - |
| | cycline | emrY | - | emrY |
| | Disinfecting agents/antiseptics | OpmH; qacH; TriA; TriB | qacH | - |
| | fosfomycin | abaF | abaF | - |
| | MLS | AmvA; AbeS; LlmA; lmrD; MacA; MacB; mefE; mphB; MuxA; MuxB; MuxC; RlmA(II) | AmvA; mphB; AbeS; RlmA(II) | AmvA |
| | Multidrug_efflux | AbeM; AcrA; AcrB; AcrE; AcrF; AcrS; AdeA; AdeB; AdeC; AdeF; AdeG; AdeH; AdeI; AdeJ; AdeK; AdeL; AdeN; AdeR; AdeS; ArlR; ArlS; ArmR; AxyX; cdeA; CmeA; CmeB; CmeC; CmeR; CpxR; efmA; efrA; efrB; emeA; emrE; emrK; EmrR; EvgA; EvgS; GadW; GadX; GolS; hmrM; H-NS; KpnE; KpnF; KpnG; KpnH; lmrP; LmrS; MarA; MdsA; MdsC; MdtA; MdtB; MdtC; MdtE; MdtF; MdtG; MdtH; MdtK; MdtM; mdtN; MepA; MepR; MexA; MexB; MexC; MexD; MexE; MexH; MexI; mexJ; mexK; MexL; mexM; MexN; MexV; MexW; MexX; MgrA; MtrA; MtrC; MtrD; OpmB; OpmD; opmE; OprJ; OprM; OprN; PmpM; RamA; SdiA; smeA; smeB; smeC; smeD; smeE; smeF; smeR; smeS; TolC | AbeM; AcrA; AcrB; AcrF; AdeA; AdeF; AdeG; AdeH; AdeI; AdeJ; AdeK; AdeL; AdeN; AdeR; AdeS; ceoA; ceoB; CpxR; CRP; EmeA; emrK; EmrR; EvgA; EvgS; GadW; GadX; H-NS; KpnF; KpnG; KpnH; MarA; MdtA; MdtB; MdtC; MdtE; MdtF; MdtG; MdtM; mdtN; MexB; MexC; MexD; MexF; MexI; mexK; MexW; MtrA; OprN; RamA; smeA; smeB; smeC; smeD; smeE; smeF; smeR; smeS; TolC | AbeM; AcrA; AcrB; AcrE; AcrF; AcrS; AdeF; AdeG; AdeI; AdeK; AdeL; ceoA; ceoB; CpxR; CRP; emrE; emrK; EmrR; EvgS; GadW; GadX; H-NS; MdtA; MdtB; MdtC; MdtE; MdtF; MdtG; MdtH; MdtM; mdtN; MexB; MexF; mexK; MexW; MgrA; MtrA; OprZ; smeB; smeC; smeE; smeF; TolC |
| | mupirocin | ileS (isoleucyl-tRNA synthetase) | - | - |
| | nitroimidazole | MsbA | MsbA | MsbA |
| | peptide | basS; arnA; bacA; BahA; basS; ICR-Mo; lipid A 4'-phosphatase; MprF; PmrC; PmrE; PmrF; YojI | PmrC; bacA; PmrE; PmrF; arnA; ICR-Mo | PmrC; bacA; PmrE; YojI; arnA; PmrF |
| | quinolone | AbaQ; EmrA; emrB; mfpA; NorA; PatA; PatB; PmrA; qacA | AbaQ; emrB; EmrA; mfpA; qacA | emrB; AbaQ |
| | rifampicin | efpA; rphB; rpoB | rpoB; rpoB2; efpA | rpoB |
| NCBI | aminoside | aminoglycoside O-phosphotransferase APH(2'')-If; aminoglycoside 6'-N-acetyltransferase; aminoglycoside O-phosphotransferase APH(3')-IIc | aminoglycoside 6-N-acetyltransferase AacA34 | - |
| | β-lactamase | class D beta-lactamase CDD-1; class A broad-spectrum beta-lactamase CfxA6 | - | - |
| | bleomycin | bleomycin binding protein BLMT; bleomycin binding protein | bleomycin binding protein BLMT | - |
| | fosfomycin | FosB family fosfomycin resistance bacillithiol transferase; fosfomycin resistance glutathione transferase FosA8 | fosfomycin resistance glutathione transferase FosA8 | - |
| | Glycopeptide/Cycloserine | VanA; VanA-Pt2; VanA-type VanS; VanB VanB-type VanR and VanS; VanC1; VanC2/3; VanC-type VanR and VanS; VanD; VanD-type VanR and VanS; VanG; VanG-Cd; VanG-Cd-type VanS; VanG-type VanR and VanS; VanH-A; VanH-B; VanH-D; VanI-type VanR; VanO-type VanR; VanT; VanT-C; VanT-Cd; VanT-G; VanU; VanW; VanW-B; VanW-G; VanX-A; VanX-B; VanX-D; VanX-E; VanX-I; VanXY-C; VanXY-G; VanXY-G2; VanY-A; VanY-B; VanY-D; VanY-G1; VanZ1; VanZ-A | VanPt-type VanS; VanX-Pt; VanO-type VanR; VanI; VanI-type VanR; VanF-type VanS; VanA-Pa; VanY-Pt; VanF-type VanR; VanZ-A; VanA-Sc; VanX-I; VanX-Sc; VanY-like; VanJ; VanX-O; VanA-type VanR; VanA-Pt | VanO-type VanR |
| | MLS | lincosamide nucleotidyltransferase Lnu(AN2); ABC-F type ribosomal protection protein Eat(A) | - | - |
| | Multidrug_efflux | multidrug efflux RND transporter permease subunit OqxB12 | - | - |
| | mupirocin | mupirocin-resistant isoleucine--tRNA ligase MupA | - | - |
| | peptide | phosphoethanolamine--lipid A transferase MCR-10.1 | phosphoethanolamine--lipid A transferase MCR-10.1 | - |
| | phenicol | - | chloramphenicol hydrolase | - |
| | rifampicin | - | rifamycin-inactivating phosphotransferase RphC | - |
| | streptothricin | streptothricin N-acetyltransferase SatA; streptothricin N-acetyltransferase Sat2 and Sat4 | - | - |
| ResFinder | MLS | mre(A) | - | - |
| | Multidrug_efflux | mdf(A); mdt(A); optrA | mdf(A) | - |
| | nitroimidazole | nimF | - | - |
| | phenicol | fexB | - | - |
| | streptothricin | str | - | - |
| ResFinderFG | aminoglycoside | Aminoglycoside N(6')-acetyltransferase type 1; Streptomycin 3''-adenylyltransferase; Bifunctional AAC/APH; 16S rRNA (guanine(1405)-N(7))-methyltransferase | Gentamicin 3-N-acetyltransferase | Bifunctional AAC/APH |
| | β-lactamase | Extended-spectrum beta-lactamase PER-1; Metallo-beta-lactamase type 2; Beta-lactamase; Beta-lactamase 3; Beta-lactamase OXA-10; putative beta-lactamase YbxI; putative peptidoglycan transpeptidase PenA; Beta-lactamase 1 | Beta-lactamase 3; Beta-lactamase OXA-10; Beta-lactamase; Extended-spectrum beta-lactamase PER-1; Metallo-beta-lactamase L1 type 3; putative beta-lactamase YbxI | Beta-lactamase |
| | cycline | Tetracycline resistance protein TetM from transposon TnFO1; Tetracycline repressor protein class A from transposon 1721; Tetracycline resistance protein TetO; Tetracycline resistance protein class C | - | - |
| | Glycopeptide/Cycloserine | D-alanine--D-alanine ligase; D-alanine--D-alanine ligase B; D-alanine--D-alanine ligase A; Vancomycin C-type resistance protein VanC; UDP-N-acetylmuramoyl-tripeptide--D-alanyl-D-alanine ligase; Vancomycin B-type resistance protein VanB; Vancomycin/teicoplanin A-type resistance protein VanA | D-alanine--D-alanine ligase; D-alanine--D-alanine ligase B; D-alanine--D-alanine ligase A | D-alanine--D-alanine ligase; D-alanine--D-alanine ligase B; D-alanine--D-alanine ligase A; Vancomycin B-type resistance protein VanB |
| | phenicol | Chloramphenicol acetyltransferase; Chloramphenicol acetyltransferase 2 | Chloramphenicol acetyltransferase | - |
| | Sulfonamide/trimethoprim | Dihydrofolate reductase; Dihydrofolate reductase type 3 | Dihydrofolate reductase type 3; Dihydrofolate reductase | Dihydrofolate reductase; Dihydrofolate reductase type 3 |
